# Noise Reduction in Resource-Coupled Multi-Module Gene Circuits through Antithetic Feedback Control

**DOI:** 10.1101/2024.05.24.595570

**Authors:** Suchana Chakravarty, Rong Zhang, Xiao-Jun Tian

## Abstract

Gene circuits within the same host cell often experience coupling, stemming from the competition for limited resources during transcriptional and translational processes. This resource competition introduces an additional layer of noise to gene expression. Here we present three multi-module antithetic control strategies: negatively competitive regulation (NCR) controller, alongside local and global controllers, aimed at reducing the gene expression noise within the context of resource competition. Through stochastic simulations and fluctuation-dissipation theorem (FDT) analysis, our findings highlight the superior performance of the NCR antithetic controller in reducing noise levels. Our research provides an effective control strategy for attenuating resource-driven noise and offers insight into the development of robust gene circuits.

## 1 Introduction

One fundamental challenge in synthetic gene circuit design is posed by resource competition, which causes unintended interplay between circuit modules [1–7]. Specifically, this interplay arises from the competition for limited cellular resources, such as RNA polymerases (RNAPs), ribosomes, and transcription factors, among the exogenous genes within the same host cell [8–10, 10–12]. This undesired coupling not only alters the deterministic behavior of synthetic gene circuits dramatically but also diminishes their robustness by inducing significant fluctuations in gene expression. For instance, resource competition manifests a linear interdependence in a simple two-gene system [9], results in non-monotonic rather than the expected monotonic dose-response curves in an activation cascade circuit [10], and leads to winner-takes-all behavior instead of the anticipated activation of two modules in cascading bistable switches circuits [12]. In addition, resource competition was found to play a dual role in gene expression noise, acting as both a novel source of noise and a regulator that constrains the level of fluctuations [13]. The uncertainties caused by context factors and noise in a gene circuit may result in circuit failure [1, 14]. This underscores the imperative for developing techniques that mitigate the repercussions of resource competition on both deterministic and stochastic behaviors to enhance the robustness of synthetic gene circuits.

One effective strategy involves leveraging orthogonal resource systems, such as orthogonal ribosomes and RNA polymerases, to establish distinct resource pools for genes [15–17]. Another widely employed approach is to incorporate negative feedback or incoherent feedforward control mechanisms into the circuit design [18–25]. These works have demonstrated the effectiveness of these mechanisms in controlling one of the single modules in the system. Interestingly, multi-module control strategies have been proposed to alleviate winner-take-all resource competition [26, 27].

Despite these advancements, effective control strategies for reducing gene expression noise within the context of resource competition are still lacking. Our previous work has initiated the investigation of the noise reduction capabilities of multi-module negative feedback control strategies [13]. Here, we proposed several antithetic controllers using different architectural forms, including single-module controller (SMC), local controller (LC), global controller (GC), and negatively competitive regulation (NCR) controller. We compared their noise reduction efficiency in the two-gene system under different levels of resource competition and fluctuation. Our finding reveals that the NCR antithetic mechanism is optimal for noise reduction within the context of resource competition. This research highlights the efficiency of the NCR mechanism in conjunction with antithetic control for enhancing noise mitigation in synthetic multi-module gene circuits.

## 2 Design and formulation of Antithetic Controllers

The two-gene circuit with two similar but independently controlled genes in the same cell is widely used to characterize gene expression noise and resource competition [9,28]. Here we use this two-gene system to study the noise reduction efficiency of several antithetic controllers. As shown in Fig. 1a, we considered the transcription of GFP and RFP genes for mRNAs, translation for GFP and RFP proteins, and degradation of both mRNAs and proteins. In addition, we considered the competition between two modules for shared transcriptional and translational resources. This resource coupling contributes to the overall protein noise as the fluctuation of one mRNA leads to the fluctuation in the available translational resource for the other mRNA [13].

**Figure 1:**
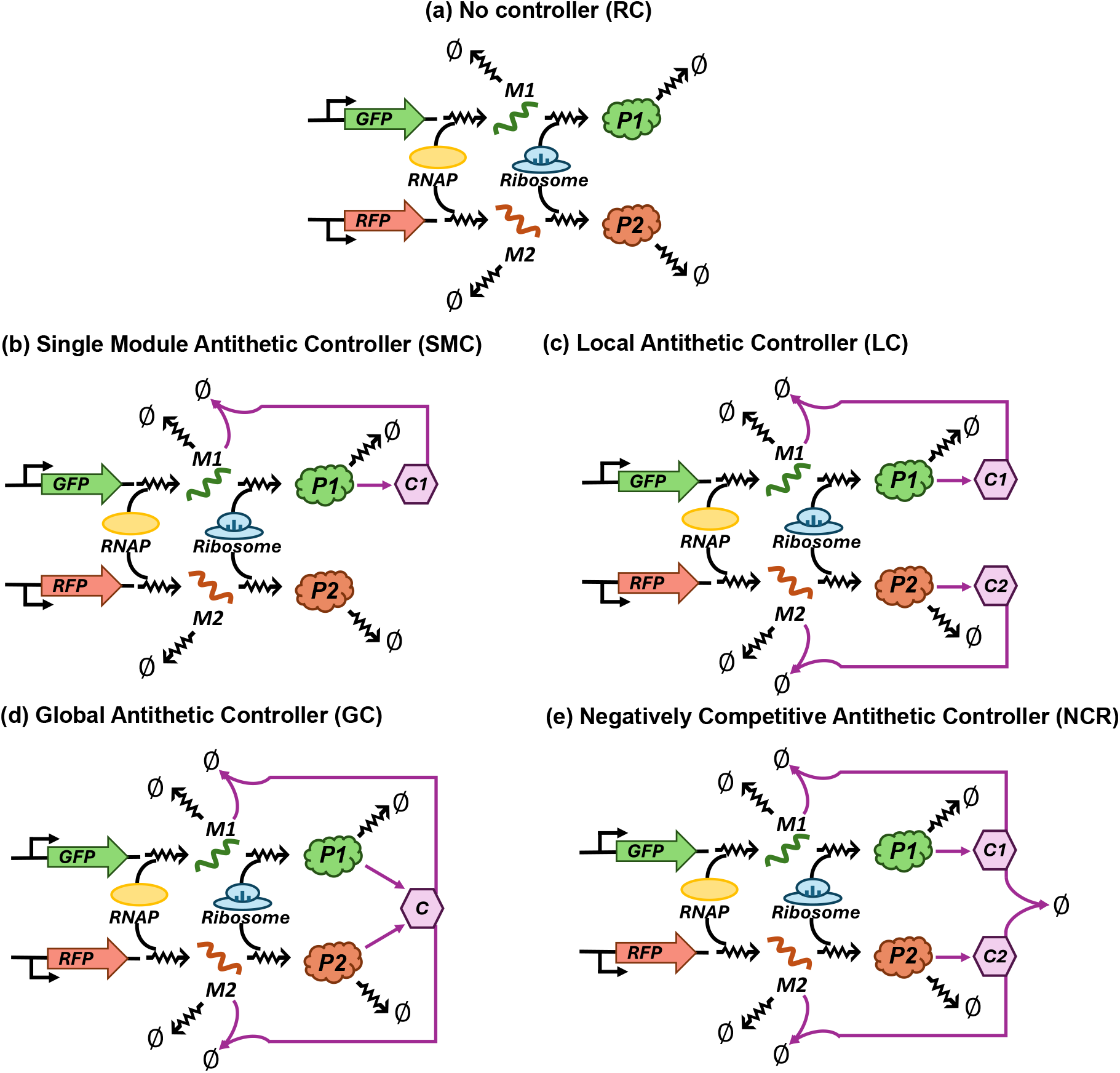
Antithetic feedback control strategies for gene expression noise reduction. (a)Diagram of the system with resource competition (RC) between GFP and RFP modules in the absence of any controller. (b) Diagram of the system with a single module antithetic negative feedback controller (SMC), where a single controller node is produced by the GFP module to facilitate the degradation of GFP mRNA through base pairing. (c) Diagram of the system with local controller (LC), where one controller node is produced by each module protein to independently regulate its own module mRNAs. (d) Diagram of the system with a global controller (GC), where a common controller node is produced by both modules to promote the degradation of both module mRNAs. (e) Diagram of the system with a negative competitive regulation module (NCR), which comprises two separate negative feedback loops similar to LC but also involves the joint degradation of the two controller nodes.

Antithetic feedback control strategy has been employed to achieve perfect adaptation in the presence of biochemical noise and environmental fluctuations [29–31]. Moreover, an antithetic controller has been used to adapt the expression of the gene of interest to variations in ribosome availability caused by resource competitors [21].

Here, we aim to test the effectiveness of antithetic controllers in reducing gene expression noise within the context of resource competition. We proposed several architectures of antithetic controllers (Fig. 1b-e) by applying the antithetic mechanism on one module or both modules of the circuit in different ways. In the case of a single-module controller (SMC, Fig. 1b), the antithetic mechanism has been implemented into the system by incorporating an antisense RNA (control node *C*1 in Fig. 1b) into the GFP module that is promoted by this module and facilitates the degradation of GFP mRNA through base pairing. Unlike the SMC model, the local controller (LC) has two distinct antisense RNAs (*C*1 and *C*2, Fig. 1c) that are promoted by two module proteins and control two module mRNAs respectively. Global controller (GC), on the other hand, employs a common antisense RNA (*C*) promoted by both modules to facilitate the degradation of both module mRNAs (Fig. 1d). The Negatively competitive regulation (NCR) controller is similar to LC but it includes an additional step where the two antisense RNAs (*C*1 and *C*2) undergo co-degradation (Fig. 1e).

We first built a general model with a set of ordinary differential equations (ODE) to describe the dynamics of mRNA (*M*_*j*_), control mRNA (*C*_*j*_), and protein (*P*_*j*_) levels for all systems with/without controllers.

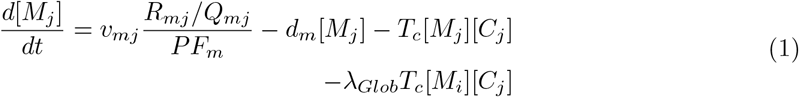

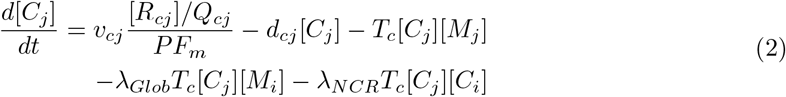

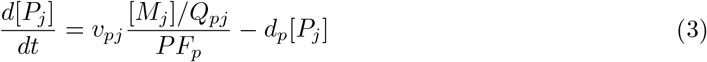

The index j here indicates module j (1 or 2), and index i signifies the regulation of molecules in module i on module j. The first terms in these are the production rates of mRNA, controller mRNA, and protein with maximum rate *v*_*mj*_, *v*_*cj*_, *v*_*pj*_ respectively. Here *PF*_*m*_ and *PF*_*p*_ define the effects of resource competition in transcription and translational processes as in our previous works [12,26,27],

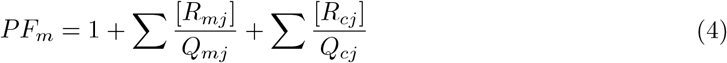

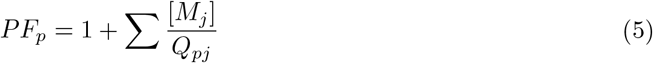

*R*_*mj*_ and *R*_*cj*_ are the fractions of active promoters for the mRNA and controller mRNA respectively. While *R*_*mj*_ is a constant, 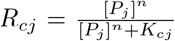 as the controller mRNA is regulated by protein, where *n* is the Hill coefficient and *K*_*cj*_ represents the protein binding affinity.

The degradation rates of mRNA, controller mRNA, and protein, are assumed to be linearly dependent on their concentration with constants *d*_*m*_, *d*_*cj*_, *d*_*p*_, respectively. In addition, the co-degradation of mRNA and its corresponding controller antisense mRNA is considered in all controllers. In the global controller, the co-degradation of mRNA and the controller antisense mRNA from the competing module is considered additionally. The co-degradation of two controller antisense mRNAs is considered in the NCR controller. Parameters *λ*_*Glob*_ and *λ*_*NCR*_ are used here to distinguish the types of the controllers. The system acts as a local controller by setting *λ*_*Glob*_ = 0 and *λ*_*NCR*_ = 0, as GC by setting *λ*_*Glob*_ = 1 and *λ*_*NCR*_ = 0, and as NCR with *λ*_*NCR*_ = 1 and *λ*_*Glob*_ = 0.

Based on this ODE model, we also built a stochastic model to simulate the stochastic trajectories of mRNA and protein levels using the Gillespie algorithm. In addition, we combined it with fluctuation-dissipation theory (FDT) to estimate the level of intrinsic noise in gene expression [32,33]. More information on these two methods can be found in our previous work [13]. Given that the mean numbers of mRNA and protein are the key determinants of the noise level of the protein, here we set their mean numbers the same for the system with/without controllers by re-scaling the transcription (*v*_*mj*_) and translation (*v*_*pj*_) rate constants throughout our research. We fixed the protein numbers at 100 while changing the mRNA numbers in the range [0, 50]. For simplicity, here we do not take into account the variation in the translational/transcriptional resources and gene copy numbers. That is, both the production rates of mRNA and controller antisense mRNA are constant in our system. The kinetic parameter values used for our simulation are, unless otherwise specified, *d*_*m*_ = 0.01, *d*_*p*_ = 0.03, *R*_*m*1_ = *R*_*m*2_ = 50, *K*_*c*1_ = *K*_*c*2_ = 250, *n* = 2, *Q*_*p*1_ = *Q*_*p*2_ = 50, *v*_*c*1_ = *v*_*c*2_ = 200, *d*_*c*1_ = *d*_*c*2_ = 1, *Q*_*m*1_ = *Q*_*m*2_ = 1, *Q*_*c*1_ = *Q*_*c*2_ = 1.

## 3 Results

### 3.1 Noise reduction by the introduction of antithetic controller mechanism

First, we examined whether the proposed antithetic controllers can reduce noise levels using stochastic simulation. The stochastic trajectories of GPF and RFP proteins, both with and without controllers, are illustrated in Fig. 2a-e. Here, the GFP and RFP mRNA numbers are fixed at 40 and 5 respectively. The small number of RFP mRNA leads to a significant fluctuation in RFP protein, consequently triggering substantial variations in GFP levels due to resource competition. This observation is evident through the anti-correlated fluctuations observed in the GFP and RFP proteins (Fig. 2a). Introducing an antithetic control strategy on either a single module (SMC) or both modules (LC, GC, NCR) has the potential to reduce the noise level, as evidenced by the decreased fluctuations observed in GFP protein trajectories and the narrower distribution of GFP expression compared to the RC case (Fig. 2f-j). Interestingly, the NCR controller exhibits the narrowest GFP distribution among all the controllers, suggesting its exceptional performance in reducing noise levels within the context of resource competition.

**Figure 2:**
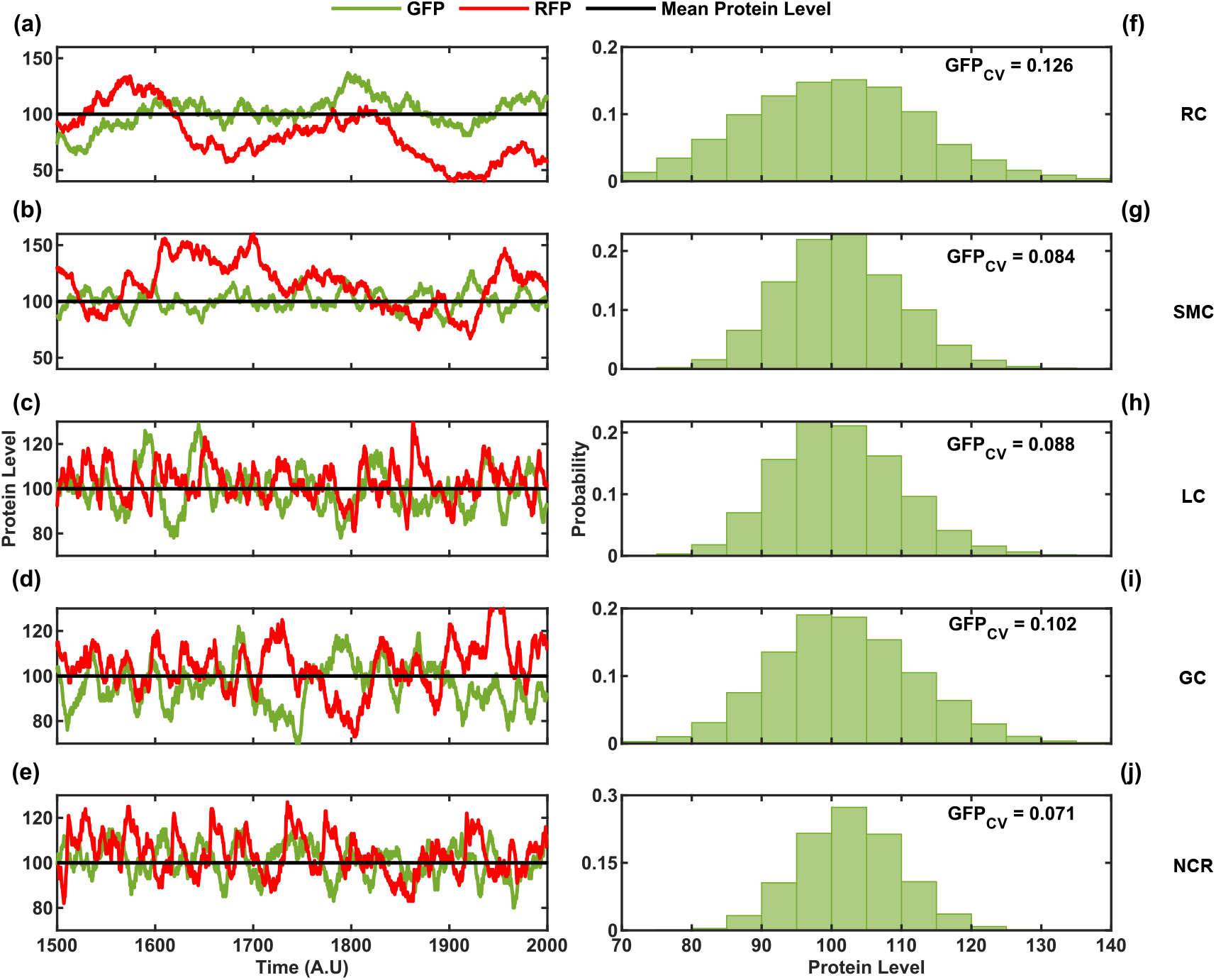
Stochastic simulation shows a noise reduction with four antithetic negative feedback control strategies. (a)-(e) Time course simulations using the Gillespie algorithm show the trajectories of GFP (green) and RFP (red) under scenarios without a controller (a), with single-module controller (b), local controller (c), global controller (d), and negatively competitive regulation controller (e), respectively. The horizontal black line indicates the mean protein level. (f)-(j) The GFP distributions for the respective models are indicated, along with their coefficient of variation. For this simulation, we assumed that the GFP and RFP mRNA copy numbers were 40 and 5, respectively, and that the GFP and RFP copy numbers were both 100. RC: no controller; SMC: single-module controller; LC: local controller; GC: global controller; NCR: negatively competitive regulation controller.

### 3.2 The performance of antithetic feedback controllers in noise reduction depends on mRNA levels

To investigate the influence of GFP and RFP mRNA copy numbers on protein noise levels, we expanded our analysis to examine the dependence of the GFP protein noise level on the quantity of RFP mRNA. Specifically, we fixed the GFP mRNA at low copy number (GFP mRNA = 5, Fig. 3a), moderate copy number (GFP mRNA = 25, Fig. 3b), and high copy number (GFP mRNA = 50, Fig. 3c). We assessed the GFP noise level under these conditions using the FDT method.

**Figure 3:**
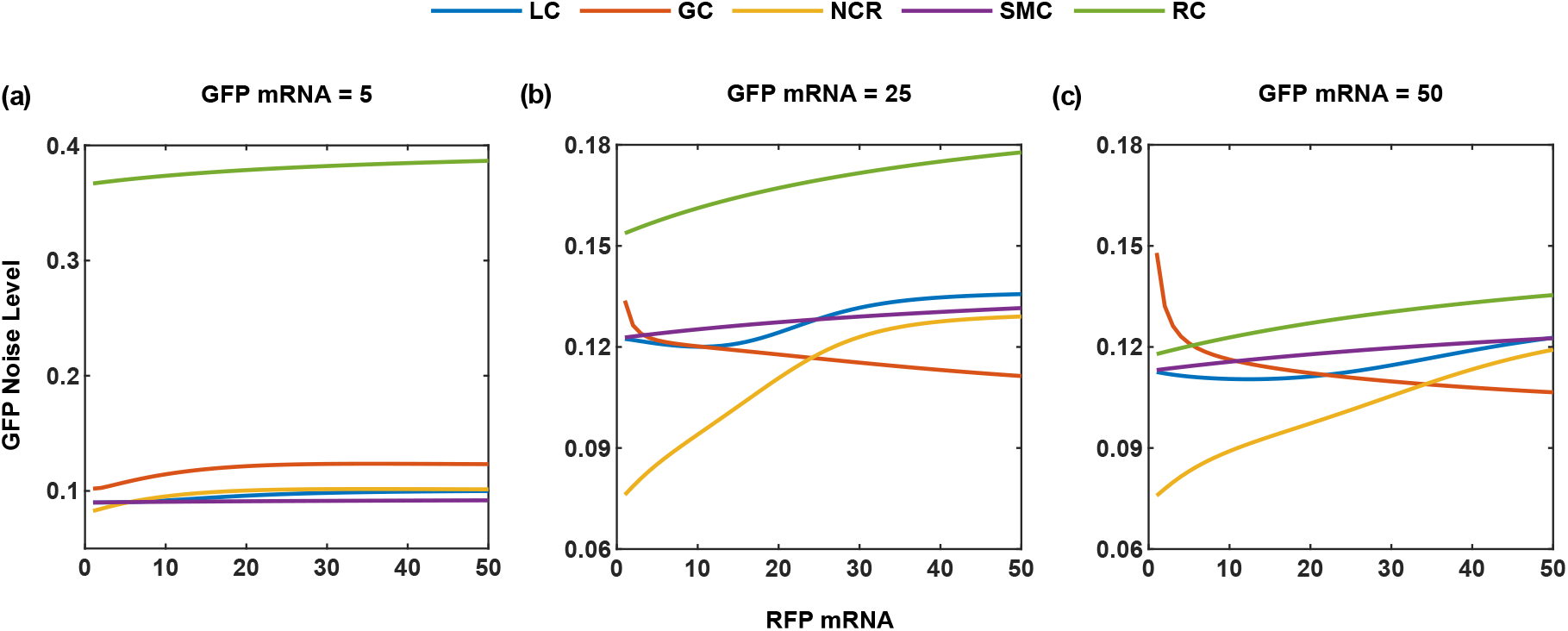
Noise reduction with four antithetic feedback controllers depends on GFP and RFP mRNA copy numbers. The GFP noise level as a function of RFP mRNA under three different scenarios where the GFP mRNA copy number is set at 5 (a), 25 (b), and 50 (c) for all four controllers. RC: no controller; SMC: single-module controller; LC: local controller; GC: global controller; NCR: negatively competitive regulation controller.

At a low copy number of GFP mRNA (Fig. 3a), a notable level of GFP noise is observed in the absence of a controller node (RC system, green line). This is attributed to minor changes in GFP mRNA resulting in substantial fluctuation in translation. Furthermore, GFP noise increases with the RFP mRNA copy number due to the resource competition-mediated coupling. Interestingly, all four antithetic controllers demonstrated significant reductions in noise levels, with the GC controller slightly less effective than others in this scenario. As we raise the copy number of GFP mRNA, the GFP noise level in the RC case decreases but still shows an increase with RFP mRNA copy number (Fig. 3b-c). The four controllers exhibit varying degrees of effectiveness in noise reduction. As shown in Fig. 3b, SMC and LC continue to demonstrate comparable levels of noise reduction. Intriguingly, the noise curves exhibit a decrease in the GC case while showing an increase in the NCR case with rising RFP mRNA copy numbers. At one critical point, the noise curves of NCR and GC intersect. Below this critical point, NCR outperforms the other controllers, while GC excels in the range of large RFP mRNA copy numbers. Additionally, it is observed that the critical point shifts to the right with a high copy number of GFP mRNA (Fig. 3c), expanding the range where the NCR works as the optimal controller. These findings suggest that GC performs optimally in regions where two competing mRNAs are comparable in abundance, whereas NCR excels in cases where the system exhibits greater asymmetry.

### 3.3 NCR antithetic controller exhibits the most effective performance in noise reduction

To further compare the performance of the four antithetic controllers in noise reduction, we systematically examined the GFP noise level in the space of GFP and RFP mRNA copy numbers. As shown in Fig. 4a, the GFP noise level significantly decreases with GFP mRNA copy number and slightly decreases with RFP mRNA copy number, consistent with Fig. 3. All four controllers demonstrated different degrees of noise reduction (Fig. 4b-e). SMC and LC demonstrate similar noise reduction capabilities across the entire space, except for a small region characterized by high GFP mRNA and low RFP mRNA copy numbers, where LC outperforms SMC slightly. Overall, both controllers exhibit a great performance with low GFP mRNA copy numbers but experience a decline as the GFP mRNA copy numbers increase. This pattern is attributed to the single control strategy in SMC and the separate control strategies in LC.

**Figure 4:**
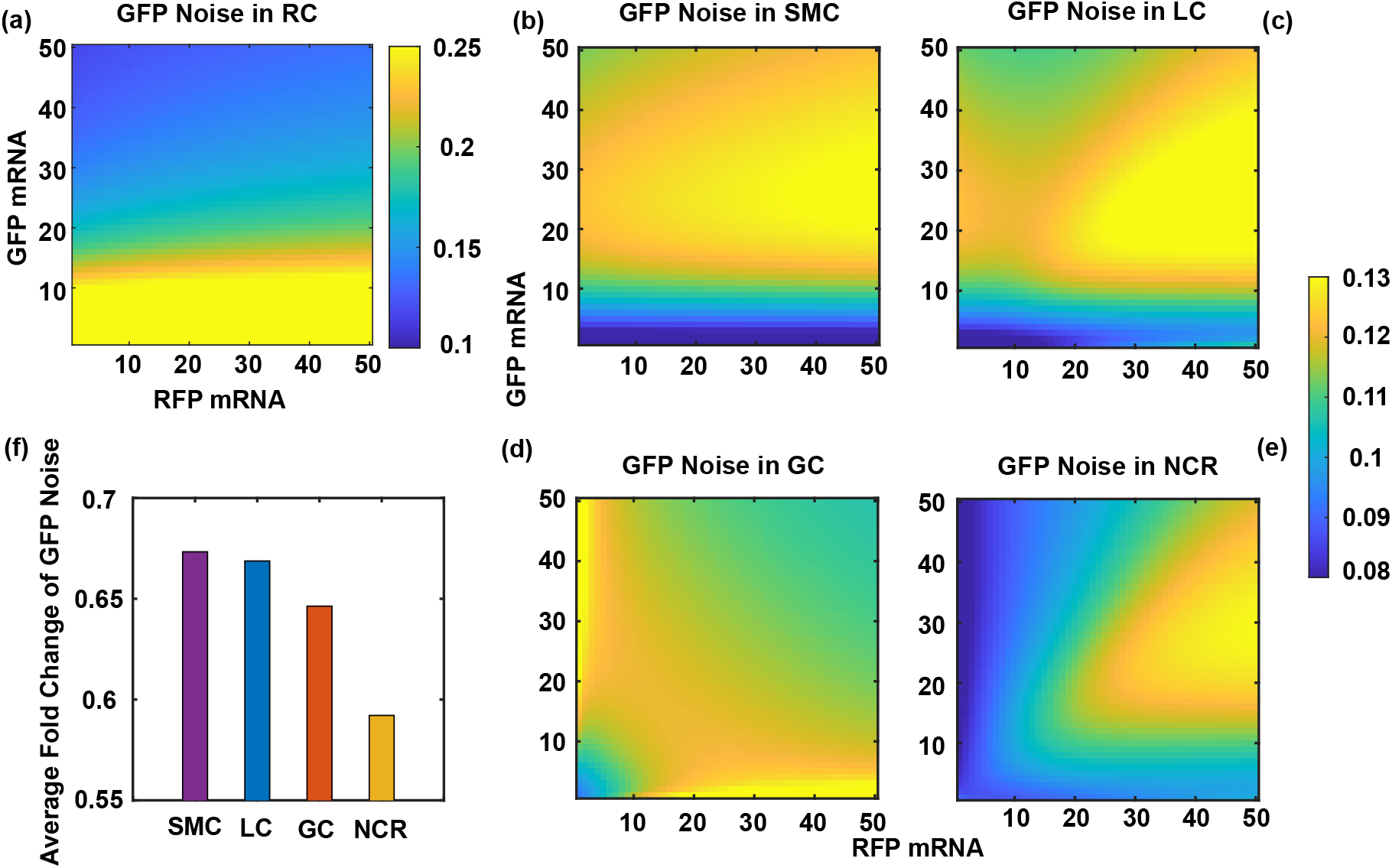
GFP protein noise within the GFP and RFP mRNAs space shows overall noise reduction performance of four antithetic feedback controllers. (a)-(e) Heatmap across GFP and RFP mRNA space depicting the level of GFP protein noise for the system without a controller (a), SMC (b), local controller (c), global controller (d), and NCR controller (e). The color bar indicates the GFP protein noise level. (f) The average fold change in GFP noise level relative to the uncontrolled system is shown for all four controllers to compare their noise reduction efficacy. RC: no controller; SMC: single-module controller; LC: local controller; GC: global controller; NCR: negatively competitive regulation controller.

Interestingly, GC and NCR show distinct patterns. GC demonstrates effectiveness in noise suppression when both GFP and RFP mRNA levels are either very low or very high (Fig. 4d), but it fares poorly in regions where GFP and RFP mRNA levels are imbalanced. In other words, GC excels at controlling noise under symmetric scenarios. The NCR system exhibits superior overall noise reduction capabilities across a wider range, particularly in asymmetric scenarios characterized by imbalanced mRNA copy numbers. The only scenario where GC outperforms NCR is when both mRNA copy numbers are high. These interesting patterns arise from the presence of cross-regulations within the NCR and GC mechanisms. In GC, the two modules may inhibit each other through the shared control node C, while in NCR, the two modules may mutually promote each other through the co-degradation of their control nodes. In the imbalanced region, the fluctuation of two mRNAs is more positively correlated due to positive cross-regulation in NCR. This positive correlation can counteract the negative correlation resulting from resource competition, leading to better performance of NCR over GC in these regions. In the region of high mRNA levels, the impact of this cross-regulation is minimal. However, GC tends to have a higher copy number of control nodes due to the shared control node and the absence of co-degradation, leading to the superior performance of GC over NCR in this region.

To assess the overall performance of these controllers, we defined noise reduction efficacy as the average fold change in the GFP noise level relative to the baseline uncontrolled RC gene circuit. Fig. 4f illustrates the noise reduction efficacy for the four controllers, conclusively demonstrating that the NCR antithetic controller achieves the most effective performance in noise reduction. These results indicate that controller architecture significantly influences their performance in noise suppression.

### 3.4 Noise reduction efficacy is determined by the controller architecture

To illustrate that controller design is crucial for effectively reducing gene expression noise, we first examined how noise reduction efficacy depends on two hyperparameters, *λ*_*Glob*_ and *λ*_*NCR*_. The parameter *λ*_*Glob*_ is specific for GC, representing the additional degradation of each module’s mRNA by the opposite control node compared to LC. The global controller system is the same as LC when *λ*_*Glob*_ = 0, and its strength increases with *λ*_*Glob*_. Similarly, The parameter *λ*_*NCR*_ is specific for the NCR controller, describing the additional co-degradation of two module’s control nodes compared to LC. The NCR system becomes the same as LC When *λ*_*NCR*_ = 0, and has a stronger controller strength with higher values of *λ*_*NCR*_.

Fig. 5a shows the dependence of the noise reduction efficacy on *λ*_*Glob*_ for the system with local, global, and NCR controllers. Given that LC and NCR systems are independent of the *λ*_*Glob*_, the average fold change in the GFP noise level remains unchanged with variations in *λ*_*Glob*_, exhibiting a flat curve. Intriguingly, the average fold change in the GFP noise level for the GC system initially decreases from the same value as LC at *λ*_*Glob*_ = 0 and then rises with increasing *λ*_*Glob*_. There exists a range of *λ*_*Glob*_ values where GC works better than LC and NCR. However, this advantage diminishes with higher values of *λ*_*Glob*_. That is, the performance of the global controller does not escalate with its controller strength. In fact, if *λ*_*Glob*_ is excessively high, the noise level might even surpass that of the no-controller scenario.

**Figure 5:**
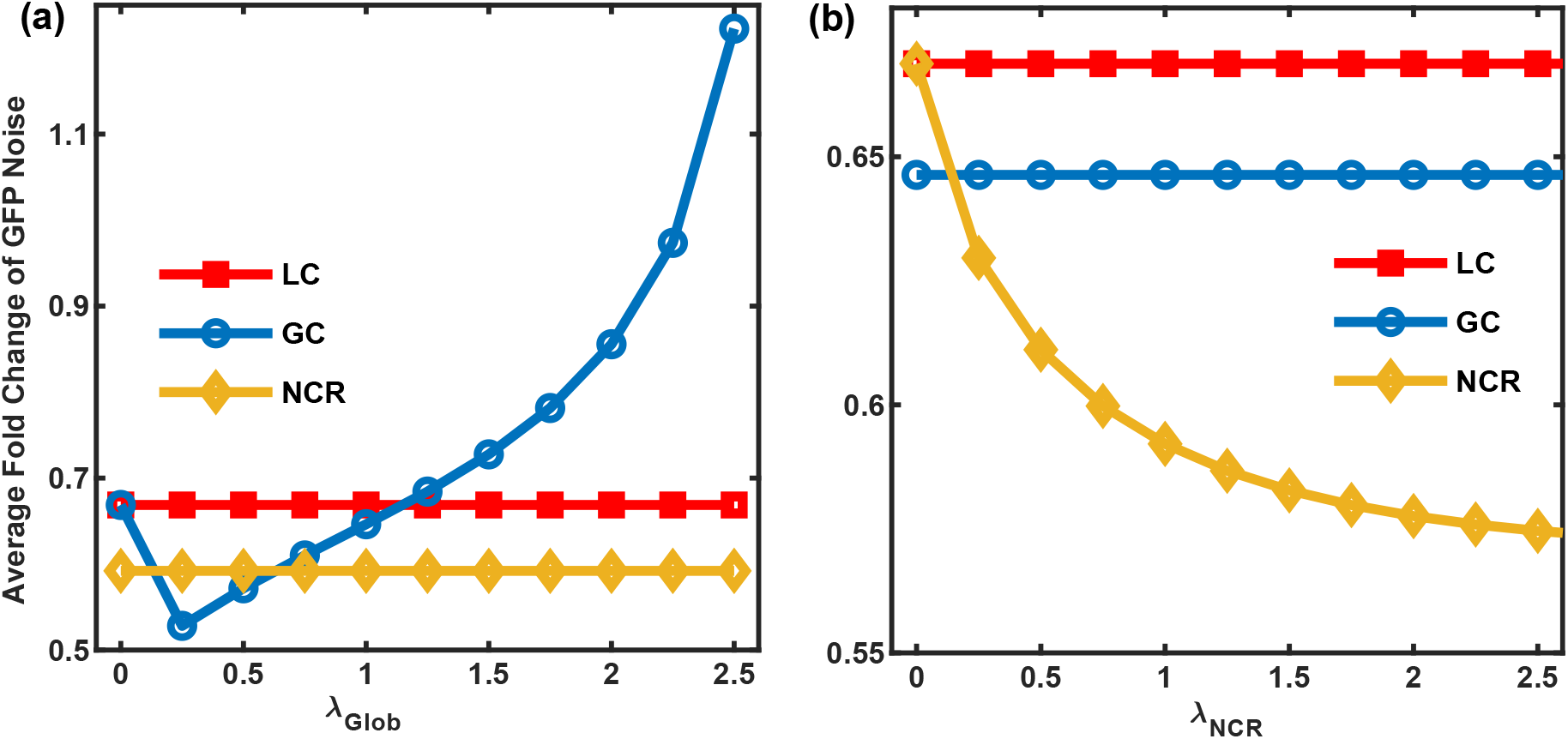
Controller architecture determines the noise reduction efficacy. The dependence of the average fold change in GFP noise level in the systems with LC, GC, and NCR controllers on parameter (a) *λ*_*Glob*_ and (b) *λ*_*NCR*_.

In contrast, increasing *λ*_*NCR*_ consistently reduces the fold change in GFP noise within the NCR system, starting from the same value as LC at *λ*_*NCR*_ = 0. Since *λ*_*NCR*_ is not involved in LC and GC systems, it does not influence the noise level in these systems (Fig. 5b). Thus, the designed co-degradation of two controller nodes offers a dependable method to minimize noise without directly affecting protein and mRNA copy numbers. These findings suggest that the NCR system is more effective and reliable in noise reduction compared to the global controller.

To further assess the effectiveness of all the controllers, we introduced extrinsic noise into the system. In order to conduct a thorough evaluation of the impacts of extrinsic noise, we generated 1000 random parameter sets with either a 20% or 50% variation to their base values. The Latin hypercube sampling method was used to ensure a representative sampling of the parameter space. Subsequently, We estimated the noise level for all the systems with each parameter configuration within the GFP and RFP mRNA space. Finally, we calculated the average fold change in GFP noise relative to the basal no-controller case. The results are shown in the scattered plots (Fig. 6) for the direct comparison of the noise reduction efficacy in the NCR system against the other three controllers. Notably, for a 20% parameter alteration, all the points fall under the diagonal line in the comparison of SMC and LC to NCR (Fig. 6a-b). Furthermore, the majority of points lie below the diagonal line in the comparison of GC to NCR (Fig. 6c), indicating that NCR outperforms the other systems across most parameter sets. With a 50% parameter variation, there is a slight increase in the proportion of points above the diagonal line in the comparison of SMC and LC to NCR (Fig. 6d-e), and 24% of points above the diagonal line for the compassion of GC to NCR (Fig. 6f). This suggests that under significant extrinsic noise, GC has some potential to outperform NCR. Nevertheless, overall, NCR remains the optimal controller even in the presence of extrinsic noise.

**Figure 6:**
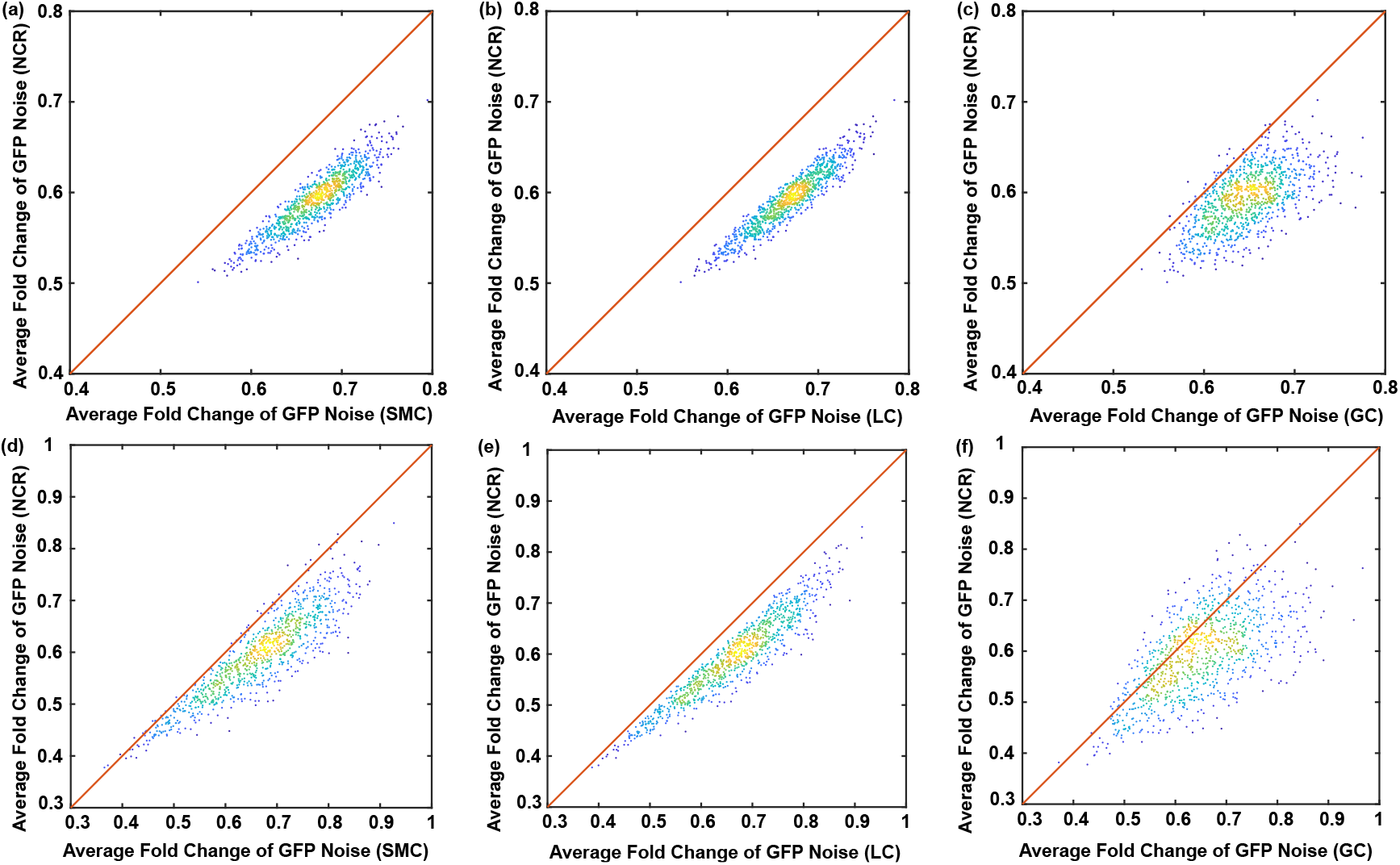
Comparison of the performance of NCR controller with respect to the other three controllers in the presence of extrinsic noise. The scatter plots illustrate the average fold change in GFP noise in the systems with an NCR controller compared to those with three antithetic controllers under 1000 random parameter sets. The value of each parameter is randomly chosen from 80% to 120% of its nominal value in (a-c), and from 50% to 150% of its nominal value in (d-f). The Latin hypercube sampling method was used to ensure a representative sampling of the parameter space.

## 4 Conclusions

Identifying the sources of noise in gene expression and controlling the level of noise are pivotal challenges in systems and synthetic biology. Traditionally, it was assumed that the cellular resources for gene expression were unlimited. However, recent findings reveal that both the translational and transcriptional resources are limited. This scarcity triggers substantial competition between genes within a synthetic gene circuit, creating indirect connections and increasing the complexity of the system. Consequently, predicting and controlling gene circuit function becomes more challenging. Additionally, fluctuations in resource availability or the biochemical reactions due to this competition introduce further unpredictability in gene expression. In this study, we introduced three distinct multi-module antithetic control mechanisms and evaluated their efficacy in controlling noise in gene expression. Through extensive simulations and systematic analysis, we demonstrated the effectiveness of these control strategies in reducing noise. Notably, we found that the negatively competitive regulation controller consistently outperforms the other controllers in noise reduction, regardless of parameter selection. Moving forward, further investigations could focus on the experimental validation of these controllers and their implementation in practical biological systems. In addition, resource competition intertwines with growth feedback, adding complexity to the system’s behavior [7, 34–38]. Exploring more sophisticated control mechanisms for managing the intricate interactions among feedback factors would be an intriguing avenue for future research.

## 5 Acknowledgement

This project was supported by the US National Science Foundation (2143229) and US National Institutes of Health (R35GM142896). The authors declare no conflict of interest.

